# Integration of customizable 3D printed mixing printheads for controlled and tuneable material and stiffness gradient fabrication with high cell viability in extrusion-based bioprinting

**DOI:** 10.1101/2025.06.22.659147

**Authors:** Florian Hofmann, Jessica Faber, Franz Moser, Alessandro Cianciosi, Nicoletta Murenu, Joachim Schenk, Katrin Heinze, Natascha Schaefer, Silvia Budday, Tomasz Jungst

## Abstract

The fabrication of native tissue-like structures with gradual transitions in material properties, cell types and growth factors remains a major challenge in biofabrication. Particularly the complex hierarchical organization, present in living tissues, has to be mimicked as close as possible for the created models to fulfil the desired function. However, the fabrication of gradual structures incorporating several materials ensuring high cell survival and subsequent unaffected tissue maturation is highly challenging. To get a step closer to the goal of generating tissue models containing controlled gradient structures, here we show a novel approach combining extrusion-based 3D bioprinting with static mixing, using self-designed Digital Light Processing (DLP) printed mixing units to fabricate defined gradual structures. Two passive mixing geometries, sinusoidal and obstacle-based, are designed and fabricated and benchmarked against commercially available static mixers. The mixing performance is assessed pixelwise using dyed alginate solutions that are extruded through the mixers using an adapted 3D printer equipped with cavity pumps. The obstacle structure exhibits the highest mixing rate compared to the sinusoidal design and commercially available static mixers. Beyond mixing efficiency, the biological compatibility of the systems is assessed by evaluating the viability of U87 and NIH3T3 cells after extrusion. Two distinct polymer solutions of differing viscosities, composed of allyl-modified gelatin (gelAGE), polyethylene glycol dithiol (PEG-2-SH), and Matrigel, are extruded through the static mixer, revealing significantly higher cell viability in the self-designed printheads compared to commercial mixers. Subsequently, graded structures mimicking the mechanical profile of native brain tissue are fabricated and mechanically characterized using nanoindentation, confirming the successful generation of continuous stiffness gradients. In summary, this study demonstrates that combining extrusion-based 3D printing with customizable mixing units enables the controlled fabrication of tissue-like gradient structures with enhanced mixing rates and cell viability. The design flexibility and rapid fabrication of DLP-printed printheads enable the adaptation to specific tissue requirements, providing a robust platform for future developments in biofabrication and tissue engineering. This work serves as foundation for the creation of increasingly complex and functional tissue models.

## Introduction

Biofabricated 3D tissue models gained increasing interest over the last decades to serve as disease models that can substitute animal experiments within pharmaceutical and cancer research. Most natural tissues, like the brain, skin, or muscles exhibit gradients with a hierarchical structure [1–3]. A gradient describes a continuous transition of materials, cells, or growth factors along an axis [4]. Since the function of living tissues is highly dependent on their hierarchical structure, the challenge within the field of biofabrication when mimicking these tissues is to resemble the spatial and temporal orientation of materials and cells as close as possible using suitable tools. If this challenge is overcome, light will be shed on the structure-function relationship and mimicry can support tissue analogues to fulfil the desired function [4–7].

In biofabrication, the biologically functional constructs are generated by automated 3D-printing processes using bioinks or biomaterial inks followed by subsequent tissue maturation steps [8, 9]. Different 3D-printing approaches like laser assisted printing [10–13], droplet-based bioprinting [13, 14], or extrusion-based bioprinting [13, 15, 16] are utilized to generate tissue analogues. In terms of bioprinted tissue analogs containing gradients, a combination of extrusion-based 3D-bioprinting and controlled fluidic mixing is one option that has been used. Printheads to address this combination were fabricated through soft photolithography [17–20], etching methods [21, 22], volumetric printing [23], hot embossing [24, 25], micro milling [26–28], or stereolithography approaches [29–31]. To generate the gradual structures, the mixing within microfluidic channels of the printheads of two or more materials can be done with active or passive mixing technologies [32, 33]. Different passive structures like those using splitting and merging of channels [34, 35], co-axial channel geometries with T- or Y-shapes with various structures as obstacles within the channel [36–38], herringbone- [39] curved sections and meander structures [40, 41], as well as chaotic mixing with static mixers [42–44] were demonstrated with different influence on mixing of the materials within those structures.

Nevertheless, the first attempts using microfluidic printheads with a Y-junction coupled to a co-axial needle extruder did not focus on mixing materials within the geometries but on extruding multi-materials within one construct, using the method to generate filaments from two materials and gradients changing materials layer-after-layer. Colosi et al. [45], for example, used microfluidic printheads to create up to 3 mm thick constructs with gradual structures by changing material composition layer-by-layer or using split filaments within one layer. Simultaneous multi-material deposition of low-viscous bioinks (blend of alginate and gelation methacryloyl (GelMA)) and 3D cell-laden constructs (human mesenchymal stem cells (hMSC) and human umbilical vein endothelial cells (HUVECs)) with bioink formulations were shown. An overall 75% cell viability after 3 days, depending on the UV-crosslinking duration, could be reached [45]. Costantini et al. [46] used a similar approach to print hydrogel fibers with muscle precursor cells (C2C12) to mimic artificial skeletal muscle tissues that were implanted in the back of immunocompromised mice. A cellular gradient of C2C12 myogenic precursors and BALB/3T3 cells within a hydrogel was created with the same layer-after- layer material change or side-by-side material extrusion within a filament as in Colosi et al. [46]. In contrast, Idaszek et al. [47] used a follow-up approach with a microfluidic printhead that was extended by a passive serpentine-based mixing structure. Hydrogel constructs (GelMA, methacrylated hyaluronic acid (HAMA) and chondroitin sulphate with 2-aminoethyl methacrylate (CS-AEMA)) with cells (hMSC and human articular chondrocytes (hACs)) and material gradients were printed for the regeneration of chondral defects [47]. The combination of the mixed hydrogels with CaCl_2_-crosslinking solution is essential for the printed constructs to retain their shape. A cell viability after extrusion around 90% was confirmed but this study only used low viscosity bioinks (1-1100 mPas) at low flow rates (0-100 µl/min). The printheads were fabricated through micro milling channels with 200 x 200 µm cross section into 5 mm thick polycarbonate sheets and subsequent hot press sealing. Adapting the mixing rates to further requirements of tissues is limited due to the diffusion-based mixing structure. It is only possible to vary the flow rates of the syringe pumps, while an adaptation of the mixing structure was not shown. Further evolvements in combination of microfluidic and extrusion-based 3D printing were demonstrated by Serex et al. [38]. Herringbone micro-mixers were added to printheads fabricated with standard photolithography and etching methods to mix colored glycerol streams with varying ratios to generate a gradual transition and the application of sheath flows and flow focusing increased the print resolution [38]. With the used printhead fabrication method a flexible and cheap adaption of the mixing structure to the requirements of different tissues is not possible due to the need of a silicon master and subsequent etching processes. Gradual structures were fabricated only by using black and white colored resins without cells or bioinks. Lavrentieva et al. [48] generated a one-layer stiffness gradient out of GelMA with different degrees of functionalization. With two syringe pumps and opposite increasing flow rates between 0 – 200 µl/min material was extruded trough a 3D-mulitjet printed microfluidic chip with a passive HC-design based mixing structure. A UV-crosslinking step fixed the gradual transition due to the different amount of homogeneous mixed functionalized material in the deposited construct. Cell viability tests with hMSC and HUVECs revealed that the used materials and the deposition process did show no significant influence on the cell viability [48]. The microfluidic chip was used to mix the materials homogeneously. An adaptation of the mixing rate was only reached due to varying flow rates. Different designs of the passive mixing structure were not tested. With the chosen broad shape of the outlet, the possibility to deposit defined 3D-gradual structures is limited. A stiffness, peptide and cell concentration 3D gradient were created with an extrusion-based 3D- printing approach by Forget et al. [49]. A bioink of native agarose was mixed with carboxylated agarose through a microfluidic printhead with a passive mixing unit inside and extruded through a nozzle. The approach consists of two syringe pumps with flow rates between 0 – 200 µl/min, a robotic arm for the deposition of materials, a temperature sensor and a cooled printing platform. Human embryonic kidney cells (HEK293) were used to generate a cell gradient and their distribution was evaluated [49]. Like described above at Lavrentieva et al. changing the mixing ratio of the materials is only possible with varying the flow rates. The setup is designed for one type of commercial mixer, a variation of the mixing structure to adapt the mixing rates is not shown.

A further option to create gradual structures are multi-material and multi-nozzle based 3D-printing approaches like the ones demonstrated by Skylar-Scott et al. [50]. They showed through printing voxelated soft matter by seamless switching between up to eight different materials to enable continuous printing. The stereolithography fabricated printheads were connected to cartridges filled with silicone inks, epoxy inks with different stiffnesses or dyed 7.5% gelatin inks. With a pressure-driven approach the constructs were printing [50]. This approach can be used to generate structures composed of different segments of materials combined in a controlled way to generate macroscopic gradients but no mixing within the channels was utilized to generate gradients within the extruded filament. Uzel et al. [51] showed another multi-material bioprinting approach where triblock copolymer, gelatin and photopolymerizable polyacrylate materials were extruded via multi-material multi-nozzle adaptive 3D-printing to mimic skin abrasions [51]. Like Skylar-Scott et al. only macroscopic gradual structures without cells were printed. Bolívar-Monsalve et al. [52] co-extruded with a syringe pump setup a myoblast (C2C12)-laden GelMA bioink along with a pristine alginate ink, using a Kenics static mixer (KSM) with three mixing elements into a CaCl_2_-bath, to chaotically mimic the skeletal-muscle like hierarchical tissue structure. The cell viability with this stretching and folding material mixing setup was around 85% up to 28 days after printing [52]. Due to the deposition of the bioink solutions into a CaCl_2_-support bath for crosslinking, no 3D structures could be generated. The variation of the mixing rate through changing the number of KSM-elements was not analyzed. Ceballos-González et al. [53]used the same multi-material bioprinting approach based on chaotic advection to generate different radial and axial micro gradients with alginate bioinks depending on the number of KSM-elements, inlets and the inlet position [53]. Macroscopic gradients were generated, but no continuous transition through mixing within the extruded filament was reached. Skylar-Scott et al. [54] presented a 3D-printing platform with multi-material multi-nozzle printheads and the option of active mixing to generate graded 3D parts. An evaluation of the mixing was done through determining the intensity of dyed materials [54]. The influence on the cell viability was not further analyzed with the shown mixing approach.

To overcome the mentioned limitations of the printhead demonstrated so far, in this work a combination of extrusion-based 3D-bioprinting and controlled fluidic mixing are used to generate gradual structures. Printheads with sub-millimeter channels to mix bioinks or hydrogels are fabricated using a stereolithography based technique to enable a fast and broad design flexibility of the mixing structures according to different requirements of the mimicked tissue, low costs, a rapid fabrication process, as well as cytocompatibility of the used materials. The accuracy of the fabrication process is evaluated by comparing the generated constructs with the concerning CAD data. An obstacle and sinusoidal passive mixing structure, are selected to mix bioink solutions containing U87 and NIH3T3 cells. With an adapted 3D-printer, the mixing ratio and quality are investigated by depositing dyed alginate solutions. To increase the resolution of the deposited structures, a nozzle adapter is integrated into the printhead to print defined structures. The cell viability of the self-designed printhead in combination with an extrusion-based setup is analyzed and compared to commercially available mixers, based on the KSM design. Gradual structures with a material and stiffness gradient are deposited with an adapted 3D-printing setup and characterized rheologically and mechanically.

## Materials and Methods

### Materials

5 ml mQ-water were pipetted in a 100 ml urine cup with screw cap. 400 mg alginate (Lot#:GQ7205601, Protanal LF 10/60 FT Sample, FMC BioPolymer, Cork, Ireland) were given in the cup and mixed with the spatula to crush large white agglomerates of alginate on the rim of the cup. Additional 5 ml mQ-water were pipetted into the cup and mixed again with the spatula. Next, the material was stored for 24 h in an incubator at 37° C. After that the 4 wt/v % solution was mixed with two tips of a spatula of green (Leaf green icing color, Wilton, Naperville, US) or red (Red icing color, Wilton, Naperville, US) food dye until the solution had a homogeneous color.

To generate the High polymer solution (5.5 % G1MM + 9.8 % PEG-2-SH + 0.33 % LAP + Matrigel), 165 mg gelatin-methacrylate (G1MM), synthesized according to an established protocol [55], and 294 mg polyethylene glycol dithiol (PEG-2-SH) (Lot#: ZZ410P048, 4ARM-SH-5000, Sigma-Aldrich, St. Louis, US) were sterilized under UV light for 20 min and then dissolved within a 5 ml cartridge with 1 ml LAP/PBS-solution, that is composed of 9 mg Lithium-Phenyl-2,4,6-trimethylbenzoylphosphinat (LAP) (Lot#: WXBD7136V, Lithium-Phenyl-2,4,6-trimethylbenzoylphosphinat, Sigma-Aldrich, St. Louis, US) that was dissolved in 3 ml PBS (pH 7,4) within an UV-lightproof cartridge. The cartridge with the G1MM/PEG-2-SH-LAP/PBS-solution is wrapped with aluminium foil to protect it from UV-light and 500 ml PBS is given to the solution. 1.5 ml of Matrigel (Lot#: 17323008, Matrigel® Matrix Basement Membrane, Corning, New York, US) is added to the polymer solution to reach a ratio of Matrigel and polymer of 1:1, resulting in a total volume of 3 ml solution. The High polymer solution is mixed through shaking manually until a homogeneous state is reached. Before use, the material is stored in a refrigerator at 5° C for 2 h. For the Low polymer solution (2.0 % G1MM + 3.6 % PEG-2-SH + 0.33 % LAP + Matrigel) the procedure is the same with different values for G1MM (60 mg) and PEG-2-SH (108 mg). To UV-crosslink the polymer solutions, a 405 nm UV-lamp (Dr. Hönle AG, Gilching, Germany) was installed at a distance of 15 cm and switched on for 20 s with a relative intensity of 60 %.

### Fabrication of DLP-printed static mixing printheads

The self-designed static mixing printheads were created with the Solid Works software (Solid Works 2022, Dassault Systèmes, Vélizy-Villacoublay, France). CAD files of every printhead and for all different channel diameters were sliced with the Prusa Slicer (Version 2.9.0, Prusa Research, Prague, Czech Republic). An adjustment of the spatial orientation and support structures for printing were added manually within the slicing software. The files were exported into a sl1s.-format suitable to the used Prusa SL1S DLP printer (Prusa Research, Prague, Czech Republic). After a calibration of the printer, which includes the adaption of the tank and the build plate, resin-based blue transparent 405 nm FotoDent® guide (Dreve ProDiMed GmbH, Unna, Germany) was put into the tank of the printer up to a filling level of 70 %. Next, the exposure time of the base layers was modified to 30 s per layer and to 8 s for all following 50 µm layers, before the printing procedure was started. After the successful fabrication of the printheads, the build plate was removed and the fixed constructs were detached from the build plate with a spatula. The support structures were cut off with a scalpel and the printheads were put into a grid of a tank filled with 2-Propanol (VWR International, Pennsylvania, US) of the curing and washing station CW1S from Prusa (Prusa Research, Prague, Czech Republic). All constructs were flushed for 4 min, cleaned with pressurized air for 1 min, rinsed again by hand with 2-Propanol to remove excess resin inside the channels of the printheads, cleaned again with pressurized air for 1 min and cured for 3 min at room temperature into the curing and washing station. Next, all static mixing chips were final crosslinked with a light-curing unit (Otoflash G171, NK-Optik GmbH, Baierbrunn, Germany) device with 4000 flashes. After that, the outlet of every printhead was abraded with a grinding machine (MetaServ 3000, Buehler, Leinfelden-Echterdingen, Germany) with an abrasive paper (P 500 WS Flex 18C Waterproof, Hermes Schleifwerkzeuge GmbH & Co. KG, Hamburg, Germany) and 250 rotations/min to generate an even outlet. The constructs were rinsed again with 2-Propanol and pressurized air for 1 min to remove the particles from the grinding process. Images of the constructs were taken with a Nikon Z6 equipped with a Nikon ED 200 mm macro-objective (Nikon, Minato, Japan). The commercially available printheads were purchased (Mischrohr 25D für eco-DUO330/450, VIEWEG GmbH Dosier- und Mischtechnik, Kranzberg, Germany).

### Evaluation of print accuracy and permeability

The evaluation of the permeability for the self-designed static mixing printheads was tested immediately after the printing and washing process. A 5 ml syringe (Becton Dickinson GmbH, Heidelberg, Germany) was filled with 2-Propanol (VWR International, Pennsylvania, US) and connected to one inlet of the printhead. The other inlet was closed by hand and it was tried to push the 2-Propanol through the channel of the chip. The same procedure was done for the second inlet. If 2-Propanol came out of the outlet for both cases, the printhead was labelled as permeable.

To verify the print accuracy and compare the results to the given CAD data, a method was developed to measure the channel diameter of the different printheads. An additional small rectangular was printed to one side of all tested printheads (Figure 1 SI b)). The bottom of the rectangular was roughened with an abrasive paper (P 500 WS Flex 18C Waterproof, Hermes Schleifwerkzeuge GmbH & Co. KG, Hamburg, Germany) to fix it with superglue (Sekundenkleber Supergel, UHU GmbH & Co KG, Bühl, Germany) to a glass plate. The diameter of the chips was measured before with a calliper. The glass plate with the fixed construct was mounted into a cutting device (EXAKT 300CL, EXAKT Advanced Technologies GmbH, Norderstedt, Germany). With a diamond saw the constructs were cut in the middle. The fixed rest of the printhead was cut off the glass plate, that both halves of the chip could be measured. If necessary, the surface of the cut printheads was abraded carefully with a sanding device (EXAKT 400CS EXAKT Advanced Technologies GmbH, Norderstedt, Germany) with the same abrasive paper as fixing the printheads at the glass plate, to remove residues from the cutting process. Next, the halved constructs were coated with a 4 nm thick platinum layer. Therefore, the samples were put into a sputter coater (Leica EM ACE600, Leica Microsystems GmbH, Wetzlar, Germany) for 45 min and the platinum layer was coated under vacuum conditions. Images of the coated halved constructs were taken with a Nikon Z6 equipped with a Nikon ED 200 mm macro-objective (Nikon, Minato, Japan). The coated samples were transferred to a tabletop scanning electron microscope (SEM) (TM3030Plus, Hitachi, Chiyoda, Japan). With the settings 15 kV, SE and an adaption of the brightness and contrast, depending on the sample, images of the coated channels could be taken. The software Fiji (Version v1.53t, GitHub, San Francisco, US) the channel diameter of the sample was measured. For this, reference points for every geometry were selected at important positions. The diameter was measured at the two inlets, the Y-junction, two times within the mixing geometry and at the outlet. The results were transferred to Excel (Microsoft Corporation, Redmond, US) and compared to the given CAD data.

**Figure 1:**
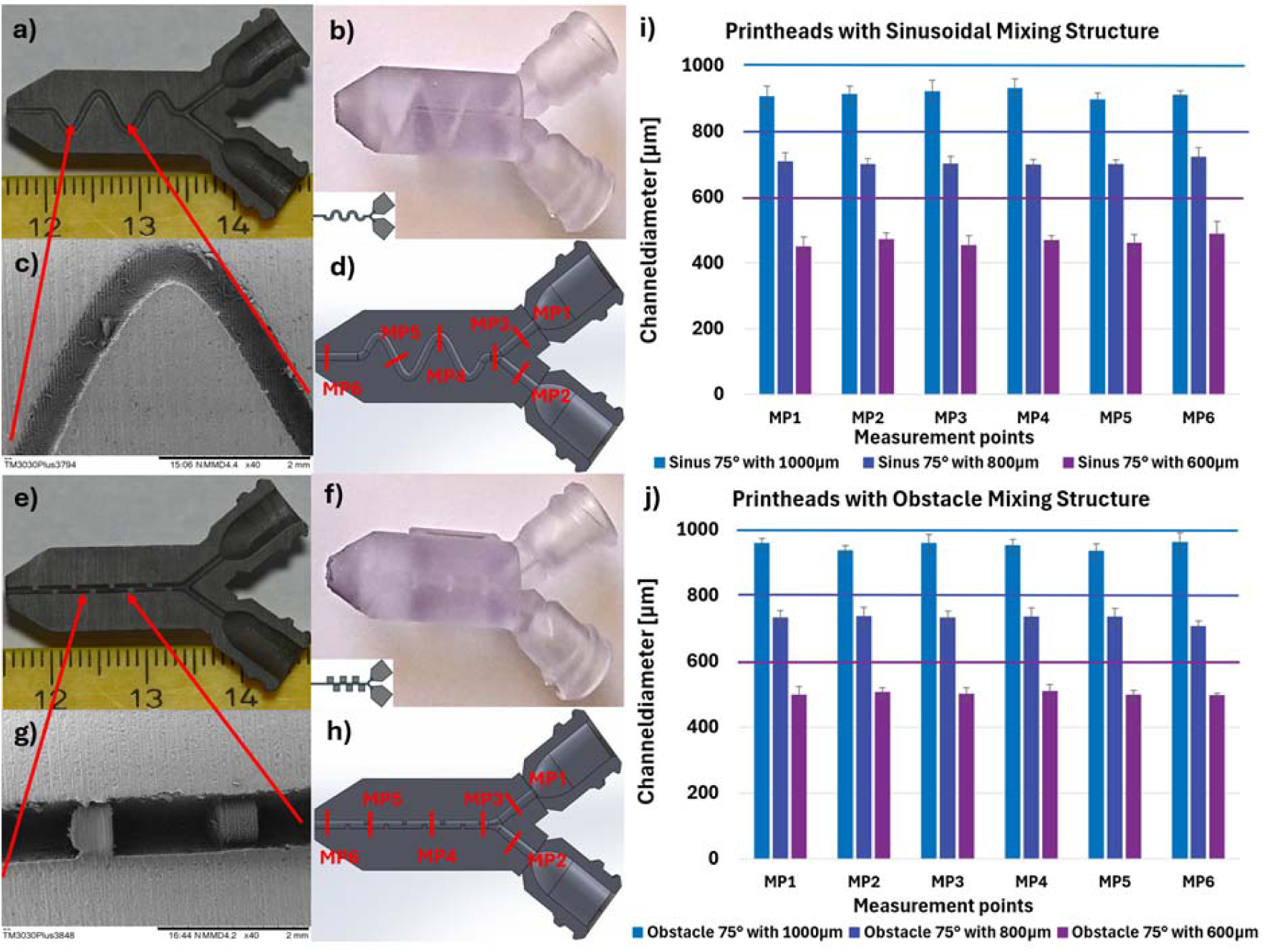
Comparison between CAD data and DLP-printed constructs. a) Printhead with a sinusoidal mixing geometry and a channel diameter of 800 µm cut in the half. b) DLP-printed printhead with a sinusoidal mixing geometry inside. c) SEM-image of the sinusoidal mixing structure. d) CAD data of a printhead with a sinusoidal mixing geometry inside with reference points. e) Printhead with an obstacle mixing geometry inside with a channel diameter of 800 µm cut in the half. f) DLP-printed printhead with an obstacle mixing geometry inside. g) SEM-image of the obstacle mixing geometry. h) CAD data of a printhead with an obstacle mixing geometry inside with reference points. i) Results of the channel diameters 1000, 800 and 600 µm at the reference points for the printheads with a sinusoidal mixing geometry inside. j) Results of the channel diameters 1000, 800 and 600 µm at the reference points for the printheads with an obstacle mixing geometry inside.

### Cell culture

U87 cells (U-87 MG, ATCC HTB-14, LGC Standards GmbH, Germany) were cultured in Dulbecco’s Minimal Essential Medium (DMEM) (41966-029, Gibco, MA, USA) supplemented with 10 % FCS (10270-106 Life Technologies, MA, US), 1 % Penicillin (50 U mL−1)/Streptomycin (50 µg mL−1) (15140-122 Life Technologies, MA, US).

NIH3T3 cells (NIH/3T3, ATCC CRL-1658, ATCC, Manassas, VA, USA) were cultivated in Dulbecco’s Modified Eagle Medium + GlutaMAX™-I [+] 4.5 g/L D-Glucose, [+] Pyruvate (DMEM) (31966-021, Gibco, MA, USA) supplemented with 10 % BCS and 1% Penicillin/Streptomycin (AC-AB-0024, Anprotec, Bruckberg, Germany). Both cell types were split twice a week.

### Evaluation of the cell viability

The viability of the U87 cells mixed with the Low and High solution was assessed after 4 days in culture. The constructs were washed with PBS (pH 7,4) and incubated for 230 min at 21 °C with 2 × 10−6 m Calcein-AM (green/living cells; Thermo Fisher Scientific, Waltham, MA, USA) and 2 × 10−6 m ethidium homodimer I (red/dead cells; Sigma-Aldrich, St. Louis, MO, USA) in PBS. Samples were imaged using an inverted Olympus IX81 microscope equipped with an Olympus FV1000 confocal laser scanning system, a FVD10 SPD spectral detector, and diode lasers of 473 nm (Alexa488) and 559 nm (Cy3) (Olympus, Tokyo, Japan). All images shown were acquired using an Olympus UPLSAPO 10× (air, numerical aperture 0.4) and were processed using ImageJ/Fiji (Version v1.53t, GitHub, San Francisco, US) and Imaris 7.7.2 (Oxford Instruments, Abingdon, UK) for the analysis of the images to quantitatively assess the number of live and dead cells. Living cells were further analysed according to cell shape. Therefore, three pictures of Calcein-AM confocal images were choses for Low and High concentration of hydrogels. Each picture was subdivided into three areas (bottom, centre, up). Images were blinded and manually counted to achieve the number of rounded versus number of elongated cells. In total n=9 per gel condition (3 pictures in total, each picture subdivided bottom, centre and up division) were analysed. Each count was performed by 3 different persons in total n=27.

The viability of the extruded U87 and NIH3T3 cells (1 × 10^6^ per ml) mixed with the Low and High solution was assessed 1 hour after printing. The constructs were washed with PBS (pH 7,4) and incubated for 45 min at 37 °C with 2 × 10−6 m Calcein-AM (green/living cells; Thermo Fisher Scientific, Waltham, MA, USA) and 2 × 10−6 m ethidium homodimer I (red/dead cells; Sigma-Aldrich, St. Louis, MO, USA) in PBS. Samples were washes again with PBS and transferred to a fluorescence microscope (Colibri 7, Carl Zeiss Microscopy Deutschland GmbH, Oberkochen, Germany) where images were taken and processed using ImageJ/Fiji (Version v1.53t, GitHub, San Francisco, US) for the analyses of the images to quantitatively assess the number of live and dead cells.

### Syringe pump setup and adapted 3D-printer

Two syringe pumps (AL-1000X, World Precision Instruments Germany GmbH, Friedberg, Germany) were used simultaneously. A 5 ml syringe (Becton Dickinson GmbH, Heidelberg, Germany) was filled with water and fixed into the holding for each syringe pump. The outlet of the syringes was connected via Luer-Lock adapters (Nordson, Westlake, US) with pneumatic tubing to two 5 ml cartridges (Optimum Kartuschen 5cc, Nordson, Westlake, US). First, the cartridges were filled with red and green dyed 4 wt/v % alginate solutions (Protanal LF 10/60 FT Sample, FMC BioPolymer, Cork, Ireland) and a plug separated the material from the water in the system. The cartridges could be connected directly to the self-designed printheads or with a special self-designed and DLP-printed adapter with the commercial mixing unit. Both syringe pumps were programmed to extrude continuously material with a resulting flow rate of 250 µl/min. Droplets were deposited into petri dishes and images were taken immediately after extrusion with a Zeiss stereomicroscope (SteREO Discovery.V20, Carl Zeiss Microscopy Deutschland GmbH, Oberkochen, Germany). The light source was chosen only from below with an exposure time of 2.6 ms. Second, the cartridges were filled with High and Low polymer solution mixed with cells. The remaining procedure stayed the same until deposition like the alginate solutions. The droplets were extruded into 96-well plates and immediately after deposition UV-crosslinked with a 405 nm UV-lamp (Dr. Hönle AG, Gilching, Germany) at a distance of 15 cm for 20 s with a relative intensity of 60 %.

A Twotrees SK1 (Twotrees, Shenzhen, China) 3D printer was adapted with a special mounting for a progressive cavity pump (PCP) (Pure Dyne kit b, ViscoTec Pumpen- u. Dosiertechnik GmbH, Töging a. Inn, Germany) to extrude material in a screw driven approach. G-codes were processed with a software (Repetier Host Version 2.2.4, Hot-World GmbH & Co. KG, Willich, Germany) and uploaded to the printer’s interface (Mainsail for Klipper, GitBook, Lyon, France). The PCP was mounted with three screws at the printer and the plastic screw was fixed at the holder designed for this purpose. First, red and green dyed 4 wt/v % alginate solutions (Protanal LF 10/60 FT Sample, FMC BioPolymer, Cork, Ireland) were filled into adapted cartridges from Puredyne and attached to the PCP. The cartridges were connected via Luer-Lock (Nordson, Westlake, US) adapters to a pneumatic tubing to further Luer-Lock adapters and the self-designed printheads or the commercial mixers. After a calibration, the samples were extruded into a petri dish at room temperature. Images of the droplets were taken immediately after extrusion with a Zeiss stereomicroscope (SteREO Discovery.V20, Carl Zeiss Microscopy Deutschland GmbH, Oberkochen, Germany). The light source was chosen only from below with an exposure time of 2.6 ms. Second, High and Low polymer solution were filled into the adapted cartridges from Puredyne and attached to the PCP. The connection stayed the same as described above, except the outlet of the self-designed printheads were connected to a 18G nozzle (Precision Tips, Nordson, Westlake, US). Constructs were deposited into a 6-well plate at room temperature and immediately after deposition UV-crosslinked with a 405 nm UV-lamp (Dr. Hönle AG, Gilching, Germany) at a distance of 15 cm for 20 s with a relative intensity of 60 %. Images were taken with a Nikon Z6 equipped with a Nikon ED 200 mm macro-objective (Nikon, Minato, Japan) of the samples within a photobox (Puluz Photo Soft Box).

### Evaluation of mixing rate and mixing quality

The images of the deposition with the syringe pump setup or the adapted 3D-printer of dyed alginate solutions were evaluated with the software Fiji (Version v1.53t, GitHub, San Francisco, US). For the determination of the mixing rate of the different printheads the images are converted into a .tif-format. Next, all areas that are not required around the sample are cut off. If necessary, air bubbles or impurities within the samples could be removed with a median-filter. With a transformation into an 8-bit-format, an image-mask could be generated. After splitting the 8-bit images into the three color channels, the red and green channels are multiplied with the mask. Next, the masked channels are divided by each other and a histogram of the final image is generated. The resulting image is divided by the maximal x-value of the histogram and the result is multiplied by 100. With the format Fire under the section Lookup Tables within Fiji, the mixing ratio of the image is pixelwise coloured displayed. A concerning calibration bar to every image could be generated. To evaluate the mixing rate an upper and lower limit for the mixing ratio are entered manually in Fiji. The software calculates the ratio of all pixels within the indicated limits for a desired area and shows the results within an Excel-file in percent. To determine the mixing quality the same conditions for the upper and lower limit of the mixing ratio have to be entered manually in the software. With a histogram the software calculates the amount of pixel for every mixing rate and displays the result within an Excel-file. The process for both evaluations is shown in Figure 3 SI.

### Rheological measurements

An Anton Paar MCR702 oscillatory rheometer (Anton Paar, Graz, Austria) with a 25 mm diameter parallel plate setup (Anton Paar, Graz, Austria) was used to measure the viscoelastic properties of the materials in torsional shear under small deformations. Before placing the materials on the measurement plate, a 405 nm UV-lamp (Dr. Hönle AG, Gilching, Germany) was mounted into the rheometer to trigger the crosslink reaction during the measurement. Next, 400 µl of hydrogel precursor was placed in the middle of the bottom plate of the device and a gap between the plates of 500 µm was chosen. After trimming the excess of material, that reached over the edge of the measurement plate, 60 µl Paraffin oil (Cas-No.: 8012-95-1, Paraffin, Sigma-Aldrich, St. Louis, US) was precisely placed around the sample to protect it from dehydration during the measurement. After that the solvent trap was pulled down to protect the material from evaporation and from ambient light that could initiate the crosslinking reaction.

With the RheoCompass software (Anton Paar, Graz, Austria) a test protocol was set up. For all following tests the temperature was set constant at 21° C. The first measurement was an amplitude sweep with a constant frequency of 10 rad / s and an increasing amplitude from values for the shear strain beginning at 0.01 %, with 10 measurement points per logarithmic decade, ending at 100 %. The storage modulus G’ and loss modulus G’’ were measured in Pa. With the amplitude sweep it was possible to determine the linear-viscoelastic-region (LVR), that characterised trough a parallel progression of G’ and G’’ to the x-axis, for following tests. The subsequent measurement was a frequency sweep with a constant amplitude of 1 %, extracted from the LVR, and decreasing frequency values beginning at 100 rad / s, with 10 measurement points per logarithmic decade, ending at 0.1 rad / s. G’ and G’’ were measured in Pa. Next, the viscosity of the materials was measured with rotational tests. The shear rate was increased from values beginning at 0.001 1 / s, with 7 measurement points per logarithmic decade, ending at 100 1 / s. Furthermore, the photo crosslinking of the two hydrogel precursor formulations was characterized by a time sweep analysis in oscillation. The parameters for shear strain and frequency were set at 1 % and 10 rad / s. After 60 s and a preparation time of 10 s to switch on the UV-lamp, the UV-lamp crosslinked the materials for 5 s with a relative intensity of 60 % trough a transparent glass bottom plate from below. After the time sweep, an amplitude and frequency sweep of the crosslinked material were measured in the same conditions of the first and second measurement. Before every test measurement the temperature was controlled and had to stay within an area of ± 0.1° C for 60 s around the test temperature of 21° C and the normal force of the rheometer was reset. After all measurements a resting step of 60 s was inserted that the material can recover from the previous test.

### Mechanical measurements

The Discovery Hybrid Rheometer HR 30 from TA Instruments (New Castle, Delaware, USA) equipped with a stainless-steel plate with a diameter of 8 mm for the upper and an anodized aluminium plate with a diameter of 40 mm for the lower specimen holder was used to measure the complex mechanical behavior of the materials in compression and tension under large deformations. The testing protocol consists of a cyclic compression-tension test with three loading cycles up to a maximum strain of 15 % in compression and 2.5 % in tension at a velocity of 40 μm/s. To improve the adhesion between the sample and the smooth specimen holders, we glued an 8 mm circular piece of fine sandpaper with a grain size of 180 µm to them. Afterwards, we added a thin layer of cyanoacrylate adhesive, Pattex superglue ultra gel (Henkel AG & Co. KGaA, Düsseldorf, Germany) to the upper specimen holder, placed the sample on a spatula and glued it to the centre of the upper geometry. Finally, we added a thin layer of the same adhesive to the lower specimen holder, inserted the transparent immersion cup, and slowly lowered the upper specimen holder with the attached sample down until full contact between the sample and the lower specimen holder could be visually confirmed. The transparent immersion cup allowed us to easily track the contact before and during testing. After a waiting time of 90 s for the adhesive to dry, the sample was immersed in a Hank’s Balanced Salt solution (HBSS) (Gibco Life Technologies, Thermo Fisher Scientific, Waltham, USA) medium bath to ensure hydration during testing. We used the advanced lower Peltier plate to perform all tests at 37 °C and mimic in vivo conditions.

### Indentation measurements

The meander structure was carefully printed on a double-sided adhesive tape which was fixed in a glass petri dish. Afterwards we filled the petri dish with Dulbeccós phosphate buffer saline (DPBS (Gibco Life Technologies, Thermo Fisher Scientific, Waltham, USA) to ensure hydration of the structure during testing. Afterwards, indentation experiments were performed at room temperature by using the Chiaro Nanoindenter (Optics11 Life, Amsterdam, The Netherlands) [56]. The machine can measure applied probe forces between 20 pN and 2 mN and displacements up to 100 µm at a resolution of 0.5 nm via an installed piezoelectric transducer. A spherical indenter with a radius of 104 µm and a cantilever stiffness of 5.0 N/m was chosen. An automated matrix scan along an indentation line with a step size of 500 µm in X direction of each horizontal strand of the meander structure was performed. During the automatic surface approach, a predefined threshold value of the cantilever deflection was reached. The displacement-controlled testing protocol consists of a constant strain rate test up to a maximum indentation depth of 8 µm and a velocity of 50 µm/s followed by a stress relaxation test at the maximum indentation depth with a velocity of 50 µm/s and a holding time of 5 s.

For data analysis, we closely followed the approaches presented in our previous studies [57, 58]. We used the solution of the axisymmetric frictionless Boussinesqs problem proposed by Sneddon [59] for the applied load F,

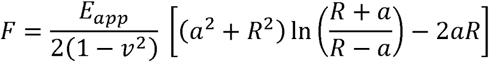

and the indentation depth h,

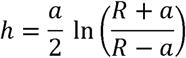

In this modified Hertz model for spherical indenters, E_app_ is the apparent modulus, which holds for deformations in the linear elastic regime, v is the Poisson’s ratio, a is the radius of the circular contact area between indenter and sample and R is the contact radius of the indenter. Under the assumption of incompressibility of hydrogels (ν = 0.5), the experimental force-displacement curves were fit to the model to determine the apparent moduli for each indentation point.

### Statistical analyses

Statistical analysis for the comparison between printed constructs and CAD data, as well as for the cell viability tests with extruded polymer solutions with cells and the evaluation of the mixing rate, was performed with SigmaPlot (Version 12.5, Systat Software GmbH, San Jose, US). Statistical significance was estimated by a µ-value lower than 0.05 (*p < 0.05, **p < 0.01, ***p < 0.001, ****p < 0.0001) using two-way ANOVA.

Statistical analysis for mechanical experiments was performed by Student’s t-test using Statistics Toolbox in MATLAB version R2024b 24.2.0.2806996 (MathWorks, USA). A µ-value lower than 0.05 was considered to be significant (*p <0.05, **p <0.01, ***p < 0.001, ****p < 0.0001).

Statistical analysis for the cell viability test with the mixed polymer solutions with cells were performed with GraphPad Prism 8.3.0 (Graphpad Software, San Diego, CA, USA) to calculate mean values, standard deviation, standard error of the mean, and values for statistical significance.

Statistical significance was estimated by a µ-value lower than 0.05 (*p < 0.05, **p < 0.01, ***p < 0.001, ****p < 0.0001) using two-way ANOVA.

## Results and Discussion

With the DLP 3D-printing approach, it was possible to generate resin-based printheads with five variable mixing structures inside the channels. A structure similar to a Y-junction was chosen for the printhead geometry, where the two inlets are on the top left and right and, after the fusion to one channel, different mixing structures were created. Variable configurations were tested for the angle between the two inlets, the shape of the outlet, the channel diameter and the adapters at the inlet for a connection to syringes. The angle between the two Luer-Lock adapters at the inlets was set to 75 °. A cone-shaped outlet with the option to add a Luer-Lock adapter for the integration of a nozzle was chosen as most suitable solution. The constructs with a sinusoidal – and an obstacle structure, representing a geometries with expected low and high mixing results [40, 41, 60], were repeatably printed and a perfusable with channel diameters as small as 500 µm were given (see Figure 1 a) SI). To verify the print accuracy and perfusability of the DLP generated printheads, a method was created to measure and compare the dimensions of the printed channels and mixing geometries to the sizes of the given CAD data (see Figure 1 b) SI).

Figure 1 shows the comparison between the printhead geometry and the CAD data for a sinusoidal (see intercept b)) and an obstacle mixing structure (displayed at section f)) for three different channel diameters of 1000, 800 and 600 µm. As shown in Figure 1 SI, the constructs were cut in the middle and sputter-coated (see segment a) and e) of Figure 1), so that it was possible to take SEM images to measure the channel diameter (shown chapter c) and g) of Figure 1). At the cutting point, it was also feasible to prove if the structure was printed correctly and if the channel is perfusable. Several reference points at distinct positions of the printhead geometry were chosen for each sample (see section d) and h) of Figure 1) at the two inlets, the Y-junction, the outlet and within the mixing structure to measure the channel diameter for comparison. The results for the sinusoidal printhead (Figure 1 section i)) and for the obstacle mixing unit (Figure 1 chapter j)) indicated an overall shrinkage for all constructs between 5 and 20 % compared to the given CAD value. Over all six reference points, the contraction for the static mixing printheads with the sinusoidal geometry is 88 ± 11 µm for a channel diameter of 1000 µm, 95 ± 8 µm for 800 µm, and 135 ± 12 µm for 600 µm. Compared to the obstacle printheads, the shrinkage for channel diameters of 1000 µm is 50 ± 11 µm, for 800 µm 71 ± 10 µm and for 600 µm 100 ± 5 µm, the results revealed for the sinusoidal structure a higher deviation than the one observed for the constructs with an obstacle structure. The values for the reference points within a channel diameter showed no significant difference for the results of the sinusoidal and the obstacle static mixing units. For both variations, the shrinkage increases with decreasing channel diameter. Straight sections like the obstacle structure have a lower shrinkage than curved parts like the sinusoidal structure. Both results can be explained since the x-y-resolution of the printer is limited, and the smaller and more complex the desired CAD structures are, the higher is the deviation. The tendency of uncured resin to stay in the channels and to close them is also higher the smaller the channel diameter becomes amplifying the deviation.

The resulting outcome shows that the used method is suitable to determine the shrinking rate and can be considered during CAD creation. The shrinkage is higher for smaller channel diameters and more complex desired structures.

The expected mixing behavior of the self-designed printheads depends on the mixing structure. The curves of the sinusoidal structure do not lead to large fluctuations in flow velocity, which means that material with higher viscosity will tend to flow side-by-side in a laminar flow through the channels and mainly diffusion induced mixing will occur through the movement [41]. For the obstacle geometry, every obstacle in the straight channel alters material flow, which leads to higher velocity differences and convective mixing can arise at these points [40, 60]. It is important to mention, that the higher the velocity differences get, the better the mixing result can be, but if working with cells, the mechanical stress to cells can also increase. This means an increase of the complexity of the mixing structure leads to an increase of mechanical stress to cells, resulting in a lower cell viability. This can be seen in the results shown in Figure 3. To evaluate the extrusion through the static mixing printheads under real conditions and confirm the expectations, it was necessary to develop a test-setup, where two materials could be extruded simultaneously through both inlets of the static mixing chips to see if mixing will occur. As materials, red and green dyed 4 wt/v % alginate solutions were chosen, because their rheology is similar to those of commonly used bioinks [45-47, 52, 53]. The tests were also performed for commercially available static mixer to compare the results to the self-designed printheads with Kenics Static Mixing (KSM) units.

With an adapted 3D-printer equipped with two cavity pumps (PCP) (see Figure 5 d)), droplets of dyed 4 wt/v % alginate solutions were extruded into a petri dish. After deposition, pictures of the droplets were taken with a stereomicroscope. To determine the mixing ratio, an image-based quantification approach as described in Figure 2 SI was established. For this, the images of the droplets were evaluated pixelwise. After image processing within Fiji, it was possible to quantify the amount of red and green color for every pixel of the droplet. The ratio is depicted by the belonging scale bar and a corresponding color pattern (see Figure 2). The range of the scale reaches from 0 to 100, where 0 means that the pixel contains only green color, accordingly, 100 means that there is only red color in the pixel spectrum. A value of 50 is representing the same intensity of red and green color in the pixel.

**Figure 2:**
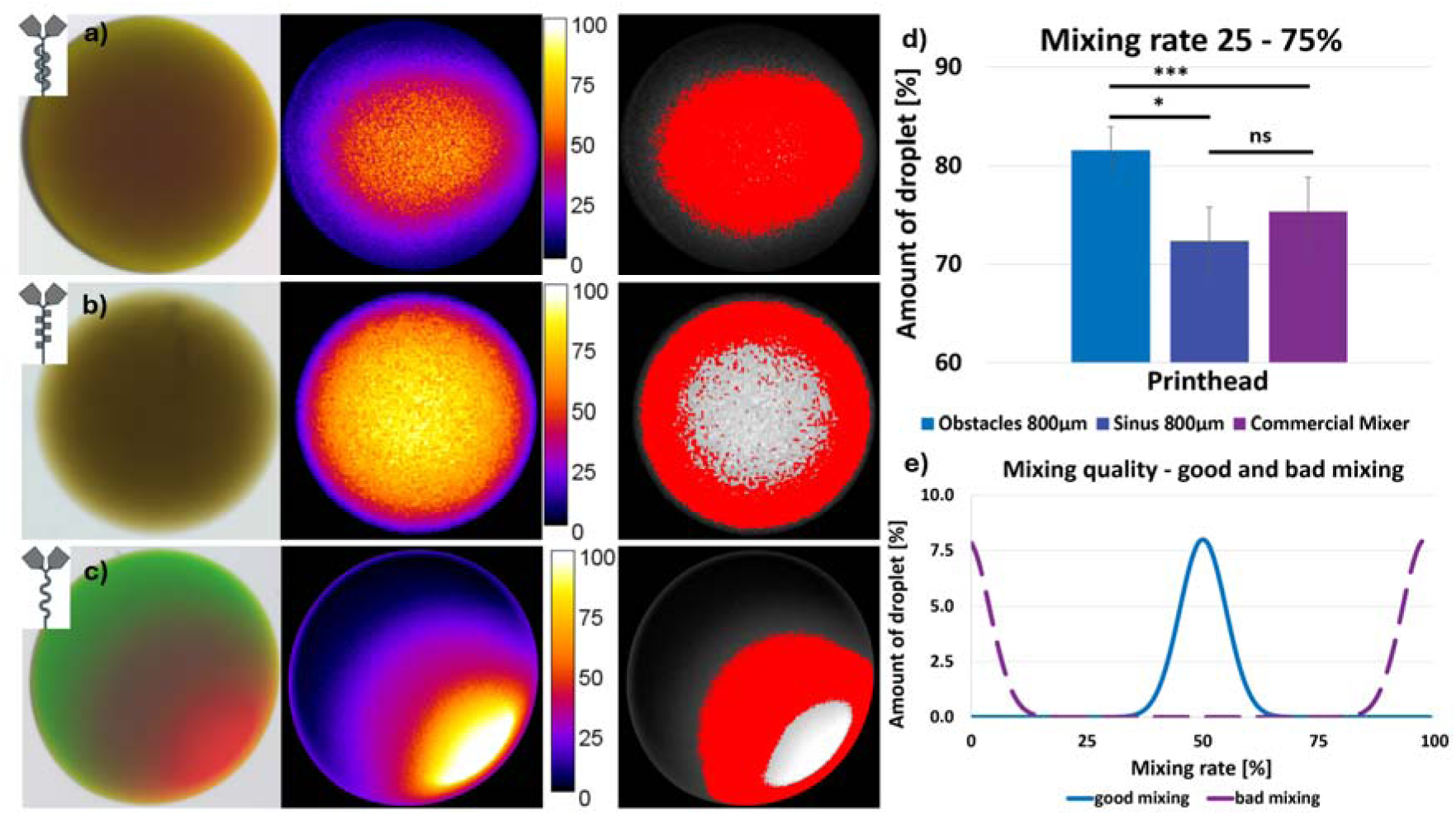
Evaluation of the mixing rate and quality of printheads with different mixing structures. a) Commercial static mixer b) Printhead with obstacle geometry c) Printhead with sinusoidal geometry; left: 4 wt/v % mixed alginate droplets with red and green food dye; middle: Evaluation of the mixing rate in 2D with a scale bar where 0 are only red and 100 are only green pixels; right: Red coloured area of the droplet where the mixing rate is between 25 and 75 %. d) Comparison of the mixing rate between 25 and 75 % for an obstacle, sinusoidal and commercial mixer. Obstacles – Sinus significant difference p < 0.001, Obstacles – Commercial significant difference p = 0.008, Sinus – Commercial no significant difference p = 0.472, significance level = 0.05. e) Theoretical mixing quality for a good and bad mixing unit.

Figure 2 shows the results for the evaluation of the mixing rate and the mixing quality of the self-designed printheads, compared to the commercially available mixer. In sections a), b) and c) of Figure 2, the following is depicted: On the left, the deposited droplet of the dyed alginate solutions. In the middle, the evaluation of the amount of red and green parts of every pixel of the image from the droplet with Fiji and a belonging scale bar. On the right, the red dyed area of mixed parts of the droplet with a certain mixing rate. The top row belongs to the commercial mixer, the results for obstacle printheads are displayed in the middle and the bottom line is part of the chips with a sinusoidal geometry. The pictures of the deposited dyed alginate droplets indicate that for the sinusoidal mixing geometry there are three parts of the droplet. From the top right to the bottom left, a green range, a mixed section between and a red area. This implies a lower mixing rate compared to the droplets of the commercial and obstacle printheads, where it is not possible to differ in red and green parts without the quantification tool. With the evaluation shown in Figure 2, it is possible to analyse the mixing behavior and compare it to commercial systems. The images on the right reveal the analysis of the pictures in the middle. The red marked area shows all pixels that have a value for the mixing ratio between 25 and 75. The bright area exhibits the mixing rates over 75 and the dark parts of the droplets offer a mixing rate below 25. To ensure that reflections on the rim of the droplets do not influence the measurements, only 90 % of the area around the centre of the droplet was considered for the evaluation. Section d) of Figure 2 shows the amount of the droplet with a mixing rate between 25 and 75 over all evaluated samples. The printheads with an obstacle structure reaches a value of 82 ± 2 % for the mixing ratio, compared to the commercial mixer with 75 ± 4 %. The sinusoidal mixers reach a mixing rate of 72 ± 3 %. These values show that the static mixing printheads with an obstacle mixing structure have significantly higher mixing ratios compared to the sinusoidal printheads and to commercial printheads. The commercial mixer shows no significant difference for the mixing ratio compared to the sinusoidal printheads.

Figure 2 e) shows different idealized outcomes of the evaluation of color distribution within the droplets as described in Figure 2 SI. If a droplet would only have values for the mixing rate of 100 and 0 of the same number, an evaluation of the whole droplet would lead to a value of 50, which indicates a complete homogeneous mixing. But this is not the case, because there are only red or green parts of the droplet, which indicates a complete inhomogeneous mixing. Due to this reason, next to the mixing rate, a further value is needed to give a statement if a mixing is good or bad. The quality of the mixing could be an indicator for this. With an image-based evaluation in Fiji, the number of every mixing rate of the droplet could be evaluated. For this, histograms of every image were created. Section e) of Figure 2 shows mixing rates between 0 and 100 within the evaluated area of the droplet for the x-axis and the number of pixels for a certain mixing rate at the y-axis. A gaussian bell curve in this diagram would be an indicator for a homogeneous mixing (blue curve). Most of the values are between a mixing ratio of 45 and 55 %. The example explained above would have a high number at 0 and 100, which is completely inhomogeneous (violet curve), but still offer a value of 50 for the mixing rate. Figure 3 SI shows the mixing quality for the commercial, obstacle and sinusoidal mixers. The results for the mixing quality for commercial and obstacle mixers are similar, both have their maxima shifted a little bit the to the right from 50 % optimal mixing rate. The results for the sinusoidal printheads show a different behavior, here are two maxima around a mixing rate of 25 and 75 %. All graphs do not exhibit an idealized homogeneous mixing, compared to the graph in section e) of Figure 2, but the results for the commercial and obstacle printheads are closer, compared to the sinusoidal ones.

**Figure 3:**
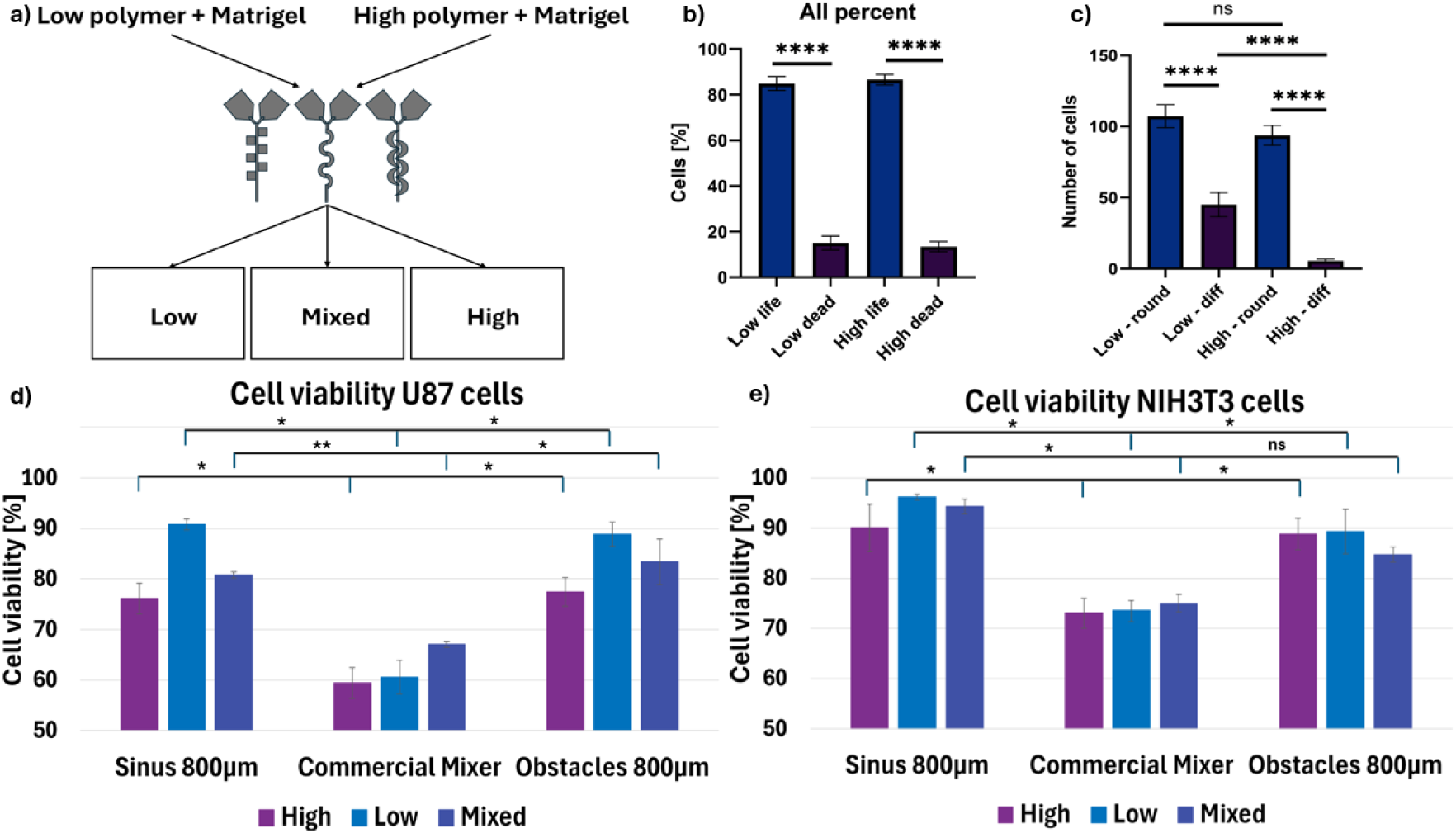
Evaluation of the cell viability for the static mixing printheads with U87 and NIH3T3 cells. a) Scheme of the setup to mix two different polymer-Matrigel solutions with the static mixing printheads to generate three diverse areas of extruded and mixed polymers. b) Amount of live and dead cells within the High and Low polymer solution. Low life – low dead and high life – high dead statistically significant with p**** < 0.0001; significance level 0.05. c) Number of cells that are round and differentiate after a period of 4 days in culture. Low round – low diff, high round – high diff and low diff – high diff statistically significant with p < 0.0001 and high life – high dead statistically significant with p**** < 0.00001; low round – high round not statistically significant with p = 0.2088; significance level 0.05.d) Cell viability of the three areas from section a) for U87-cells one hour after extrusion for printheads with a sinusoidal and obstacle geometry inside compared to the commercial mixer. Comparison of all High, Low and Mixed conditions with p*<0.001 and p**=0.001 statistically significant different; significance level 0.05. e) Cell viability of the three areas from section a) for NIH3T3-cells one hour after extrusion for printheads with a sinusoidal and obstacle geometry inside compared to the commercial mixer. Comparison of all High, Low and Mixed conditions with p*<0.001 statistically significant different and ns= 0.051 statistically not significant different; significance level 0.05.

The evaluation of the mixing rate and the mixing quality reveals that the static mixing printheads with the obstacle mixing structure have a significantly higher mixing ratio compared to commercial and sinusoidal mixers. All tested printheads do not reveal an ideal homogeneous mixing behavior, but the mixing quality of the obstacle and commercial mixer is higher compared to the sinusoidal printhead.

After the evaluation of the mixing rate and quality for the different printheads, the cell viability after extrusion of material through the mixing units was evaluated. The cell viability of the commercial mixer was compared to the printheads with the obstacle and sinusoidal mixing geometry.

To evaluate this, a test setup was developed. A scheme is displayed in section a) of Figure 3. Two polymer solutions, called „low“ and „high“, composed of varying amount of gelAGE and PEG-2-SH, including LAP as photoinitiator. These components were dissolved in a solution of Matrigel and PBS, in a ratio of 1:1 and finally cells were added under sterile conditions. Both polymer solutions were filled into cartridges, which were wrapped in aluminium foil to protect them from UV-light that no crosslinking step is initiated. The cartridges were connected with tubing and Luer-Lock adapters to a static mixing printhead or the commercial mixer. Two syringe pumps, that were connected to the cartridges also via Luer-Lock adapters with water-filled tubing extrude both polymer solutions with a flow rate of 250 µl/min into a 96-well plate. Three different bioinks termed “low”, “high” and “mixed”, were extruded. “Low” means that only material from the cartridge with the lower polymer content is pushed through the chip with a flow rate of 250 µl/min. The other syringe pump produced only a little pressure via extrusion to ensure that the material within the static mixing printhead flows in the right direction and pushing back of materials from one compartment to the other is avoided. “High” means that the material with high polymer content is extruded with a flow rate of 250 µl/min and the syringe pump with the low polymer composition was extruded at exceptionally low rates. For “mixed”, material is extruded with the help of both syringe pumps, with a resulting flow rate of 250 µl/min. After extrusion of three samples for each condition per printhead with a sinusoidal and an obstacle geometry inside and with a commercial mixer, the samples were crosslinked with a 405 nm UV-LED lamp for 20 s from a distance of 15 cm and the samples were placed in cell culture media for one hour. After a live-dead staining, images were taken with a fluorescence microscope and the cell viability was evaluated.

Sections b) and c) of Figure 3 show the cell viability and cell morphology for U87 cells only for the material without the printing process. For the “high” composition, a cell viability of 87 ± 2 % could be reached. The low condition showed with 85 ± 3 % comparable results and no significant differences. The number of round and elongated cells were for the “low” condition higher than for the solution with the “high” polymer content.

The graphs at Figure 3 d) and e) show the comparison for the cell viability of the extrusion with the syringe pumps one hour after printing for U87 and NIH3T3 cells for the three conditions with the sinusoidal, obstacle and commercial printhead. The sinusoidal printhead showed the best viability for both cell types (U87 cells: Low: 90.8 ± 1.0 %, High: 76.2 ± 3.0 %, Mixed: 80.8 ± 0.8 %; NIH3T3 cells: Low: 96.2 ± 0.7 %, High: 90.0 ± 4.7 %, Mixed: 94.4 ± 1.4 %). The obstacle printhead reached little lower viability compared to the sinusoidal mixing unit (U87 cells: Low: 88.9 ± 2.4 %, High: 77.5 ± 2.8 %, Mixed: 83.4 ± 4.5 %; NIH3T3 cells: Low: 89.3 ± 4.4 %, High: 88.8 ± 3.2 %, Mixed: 84.9 ± 1.5 %). The lowest cell viability was detected for the commercial mixer in this comparison for both cell types (U87 cells: Low: 60.6 ± 3.3 %, High: 59.4 ± 3.0 %, Mixed: 67.1 ± 0.6 %; NIH3T3 cells: Low: 73.5 ± 2.2 %, High: 73.0 ± 3.0 %, Mixed: 75.0 ± 1.8 %). These results show that the self-designed printheads have, with one exception, a significantly better cell viability, compared to commercially available mixers. For the sinusoidal printheads, the “low” condition had for both cell types the highest cell viability, the mixed section had a lower value and the “high” condition had the lowest cell viability.

These results show that the self-designed static mixing printheads enable dispensing material with a higher cell survival compared to commercially available mixers.

To assess the complex mechanical properties of the materials used for the cell viability experiments, compression tests, tension tests, and rheological measurements were performed. “Low” and “high” polymer solutions were characterized.

Figure 4 a) shows storage and loss moduli, G’ and G’’, as a function of time in a logarithmic presentation. After 65 s, the UV lamp was turned on and an increase of G’ and G’’ was the consequence. This can be explained due to the crosslinking of the polymer chains. After UV-crosslinking, all conditions have a higher value for G’, that is now also higher than G’’ (indicating the transition from a fluid to a solid). Figure 4 b) shows the viscosity of the two materials as a function of the shear rate in a logarithmic way. For small shear rate values of up to 0.01 1/s, the high polymer content solution shows a higher viscosity than the low one. Both materials showed a similar shear-thinning behavior, i.e., with increasing shear rate the viscosity decreases. Shear thinning is important when extruding the materials through a nozzle as it makes dispensing easier and can help protecting cells from mechanical stress [7].

**Figure 4:**
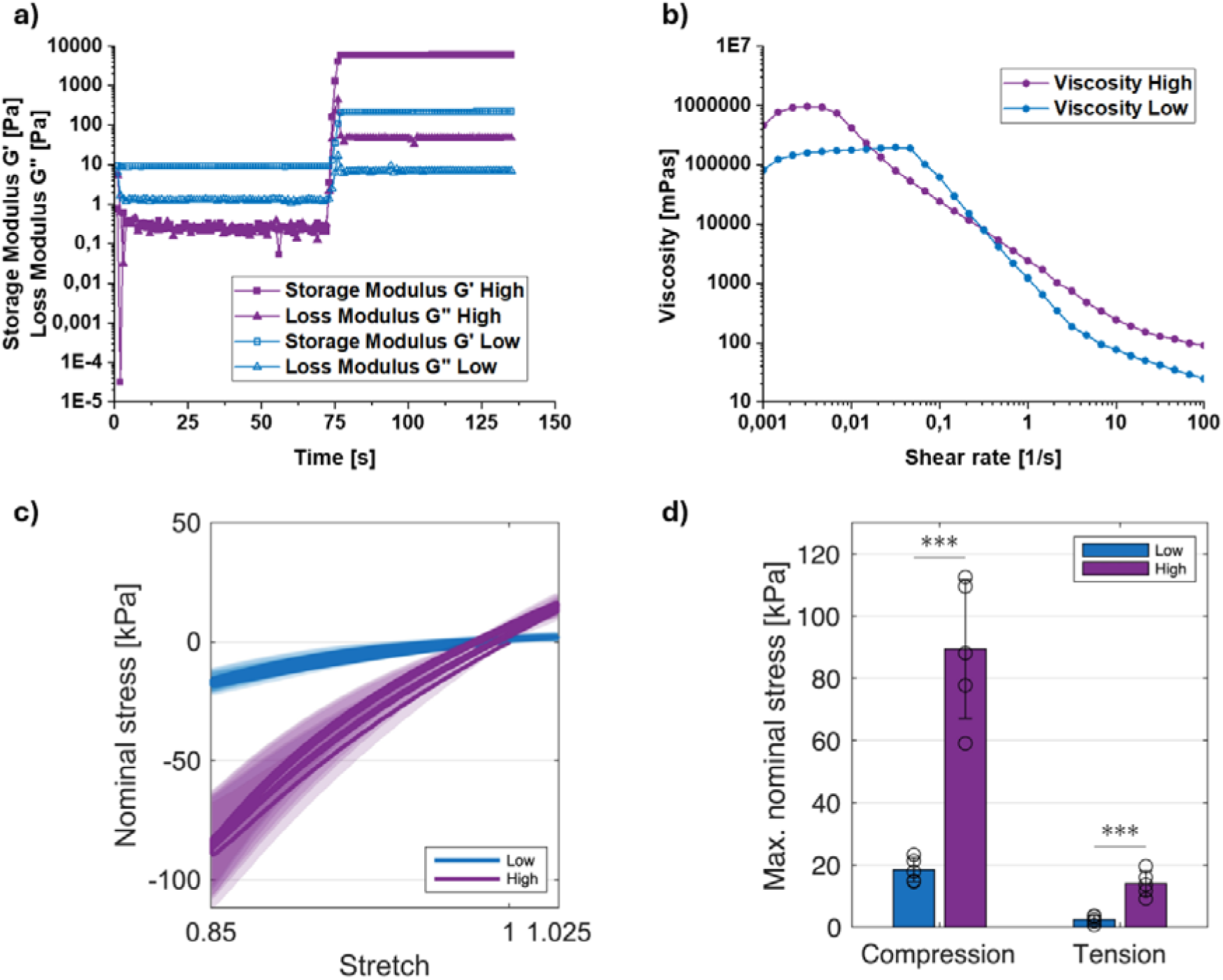
Rheological and mechanical characterisation of different polymer solutions. a) Time sweep with UV-crosslinking of polymer solutions with a low and a high polymer content. b) Viscosity sweep of polymer solutions with a low and a high polymer content. c) Average maximum nominal stresses in compression up to 15% strain and tension up to 2.5% strain of UV-crosslinked gelAGE hydrogels with a low and a high polymer content mixed Matrigel. Significances were calculated using Student’s t-test. d) Cyclic loading behavior of UV-crosslinked gelAGE hydrogels with a low and a high polymer content mixed Matrigel during cyclic compression-tension up to a maximum strain of 15 % in compression and 2.5% in tension. Significance value: ***p < 0.001.

Figure 4 c) shows the large-strain mechanical behaviour of the UV-crosslinked gels with high (n=5) and low (n=5) polymer content in compression and tension. Figure 4 d) plots the corresponding maximum stresses. Both gels only show minor viscoelastic effects during cyclic loading, i.e., hysteresis and conditioning behavior. The maximum stresses in compression and tension (Figure 4 d) indicate that the gel with high polymer content is approximately five times stiffer than the one with low polymer content under large deformations of up to 15% compressive strain.

The rheological and mechanical measurements reveal that both variants are shear thinning and that, after light induced crosslinking, the high polymer content gel has a significantly higher stiffness in shear, tension, and compression compared to the low polymer content gel.

Figure 5 a) shows the adapted 3D-printer with two PCP printheads that can be connected to the mixing units tested before to deposit gradient structures by varying the material flow in relation to the movement of the printhead. The two printheads were mounted on a special adapter and two cartridges with the mixture of a “high” and “low” polymer solution were connected via tubing and Luer-Lock adapters to the printheads. Two plugs sealed the material in the cartridges from the outside. With the PCP printheads, a constant material flow was achieved. With the aid of a G-Code, it was possible to regulate the movement of the 3D-printer to the speed of the alternating twist of the screw of every printhead that leads to material extrusion and different structures could be deposited with various ratios between the two printheads.

**Figure 5:**
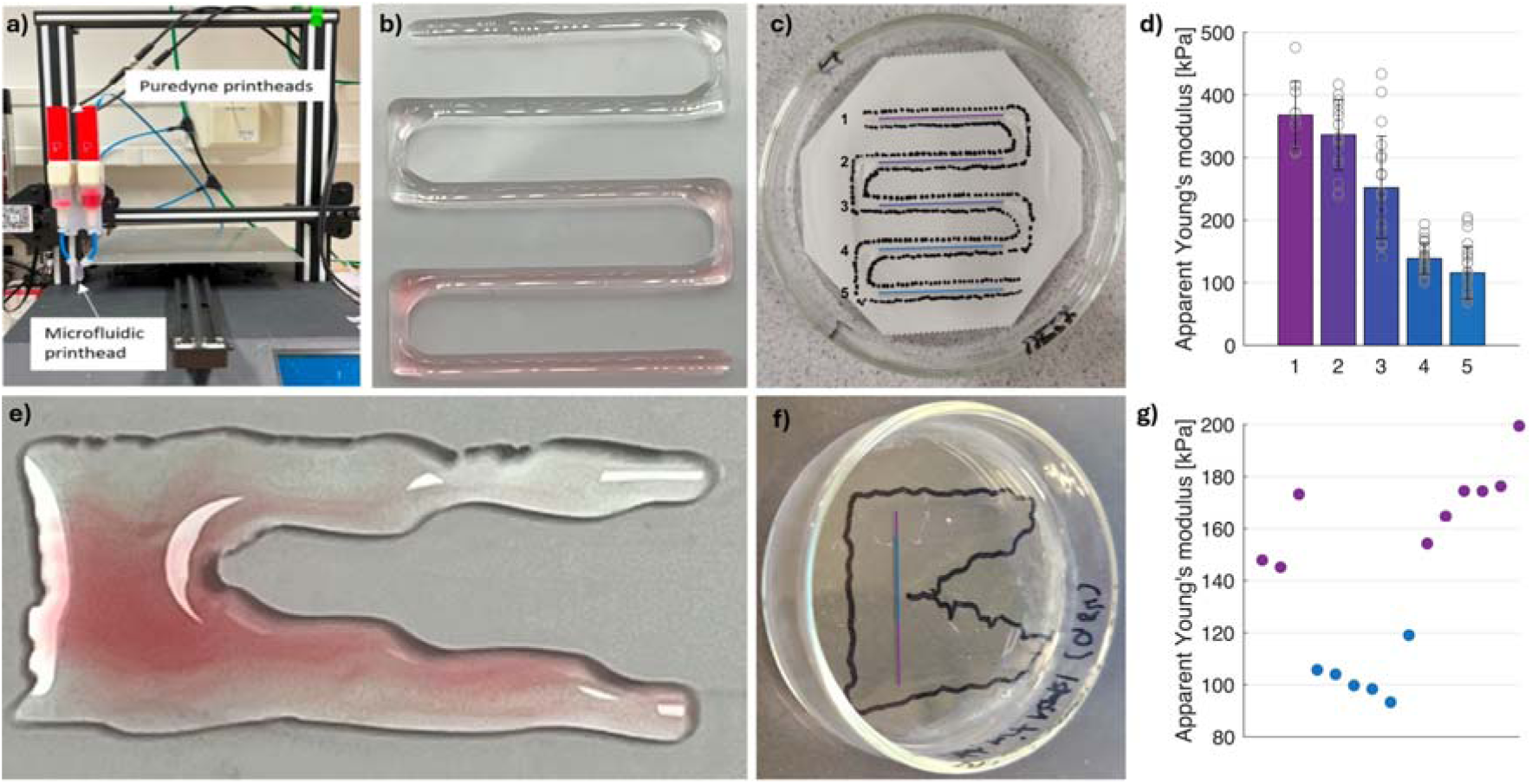
Gradient printing and mechanical evaluation. a) Adapted 3D-printing setup for extruding and mixing materials with Puredyne printheads and static mixing chips with different mixing geometries inside. b) Extrusion of meander-shaped dyed 4 wt/v % alginate strands. c) Extrusion of polymer solutions with from a high polymer content to a low polymer content inside a petri dish. d) (c.f. Suppl. Material Figure 5). e) Extrusion of native tissue-like shaped dyed 4 wt/v % alginate strands. f) Extrusion of a gradual native tissue-like structure with polymer solutions with a high and low polymer content. g) Mechanical evaluation of the line from Fig. 5f using nanoindentation.

The first print is shown in Figure 5 b), where dyed 4 wt/v % alginate strands were deposited to test the suitability of the deposition. A z-shaped structure was programmed and a transition from light green to red strands was deposited with an obstacle mixing unit in combination with a 0.84 mm diameter nozzle. The nozzle was mounted to the mixer via a self-designed Luer-Lock adapter that was printed via DLP on the tip of the printheads. Pictures were taken with a microscope immediately after printing to avoid drying of the samples. A gradient was created, starting with 100 % green dyed alginate in the first horizontal row and 0 % red dyed alginate reducing the green one by 25 % per horizontal row and increasing the red one for the same value per row to maintain a constant total material flow. The vertical rows serve only as delay distance to switch the mixing ratio of the printhead and to have time until the new ratio is extruded. Like this, a gradient was printable.

The same graded structure, starting with the high polymer solution and going to the low polymer concentration, was deposited into a petri dish, as depicted in Figure 5 c). The five colored lines indicate points where indentation measurements were performed. The results of the measurements are displayed at Figure 5 d). Line number 1, the violet line, contains only high polymer solution, resulting in the highest apparent Young’s modulus. With each horizontal line, the value for the apparent Young’s modulus is reducing. The mechanical measurements confirm the generation of the desired gradient.

Figure 5 e) displays a more complex gradual heterogeneous structure with a continuous change of material. The structure is deposited through dyed 4 wt/v % alginate strands like in section a). Figure 5 f) displays the same structure with the High and Low polymer solutions. Figure 5 g) confirms spatial changes in the local mechanical properties of the complex gradual heterogeneous structure (Figure 5 f) using nanoindentation.

## Conclusion

In this study, a novel combination of extrusion-based 3D-printing and static mixing with self-designed DLP-printed mixing units was developed to generate gradual structures. As most of living tissues exhibit gradients with a hierarchical structure and their function highly depends on this organization of materials, cells or growth factors, biofabrication needs to mimic these tissues as close as possible. Printheads with sinusoidal and obstacle passive mixing structures were created and their dimensions were compared to the CAD data for channel diameters between 600 and 1000 µm. An increasing shrinkage from 5 to 20 %, depending on decreasing channel diameter and increasing complexity of the mixing structure were the outcome. The mixing rate of the printheads were evaluated pixelwise with Fiji of droplets created from dyed mixed 4 wt/v % alginate solutions with an adapted 3D-printer with cavity pumps and compared to commercially available mixers based on the KSM design. The results revealed that the obstacle structure reached higher mixing rates (82 ± 2 %) compared to the sinusoidal structure (72 ± 3 %) and the commercial mixers (75 ± 4 %). Besides the mixing rate, another important parameter is the mixing quality, which was assessed to determine how homogeneous the mixing was. In addition, the cell viability after mixing and extrusion of the self-designed printheads was evaluated and compared to the commercially available mixers. Two polymer solutions (High and Low) mixed with Matrigel and U87 or NIH3T3 cells were extruded into 96 well plates. The results showed that the cell viability of the self-designed printheads is higher compared to the commercial mixers. A rheological and mechanical evaluation of both solutions confirmed corresponding differences. Finally, a meander-shaped gradual structure and a stiffness transition from stiffer to softer matter were deposited based on the high and low polymer solutions without cells. The created constructs were evaluated mechanically by nanoindentation to confirm the gradual property changes. The combination of extrusion-based 3D- printing and microfluidic mixing enabled the controlled mixing of materials with different viscosities and with or without cells to generate gradual transitions in material properties, while achieving higher cell viability and mixing rate compared to commercially available mixers. Due to its high design flexibility and rapid fabrication process of the self-designed printheads, it is possible to adapt the mixing structure to the requirements of different native tissues. This and the promising outcomes of this study could pave the way for further investigation of the fabrication of more complex gradual structures and tissue models.

## Supporting Information

**Figure 1:**
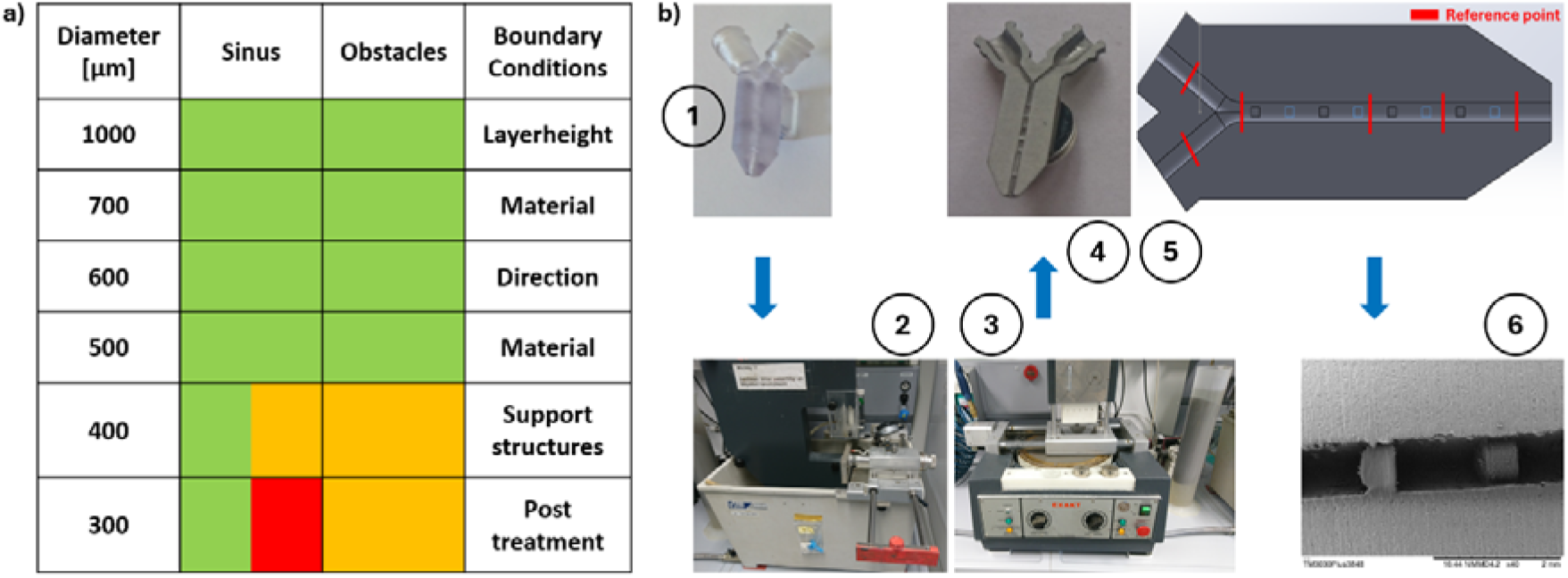
Permeability of the self-designed printheads and method for the comparison between CAD data and DLP-printed constructs. a) Table with different DLP-printed channel diameters for a sinusoidal and obstacle mixing structure. Green color represents a print- and permeable construct, red color a non-print- and non-permeable construct and orange color represents the limit where only some constructs are print-and permeable. The boundary conditions are listed on the right and are set as constant for all constructs. b) Method for the comparison between CAD data and DLP-printed constructs. 1) Printhead with an obstacle mixing geometry inside and the additional rectangular on the right side of the printhead. 2) Cutting device to half the printheads exactly in the middle. 3) Sanding device for post treatment of the surface of the halved printheads if necessary. 4) Coated halved printhead. 5) Election of distinctive reference points of the respective mixing geometry. 6) SEM image at the reference point for the evaluation of the channel diameter with Fiji.

**Figure 2:**
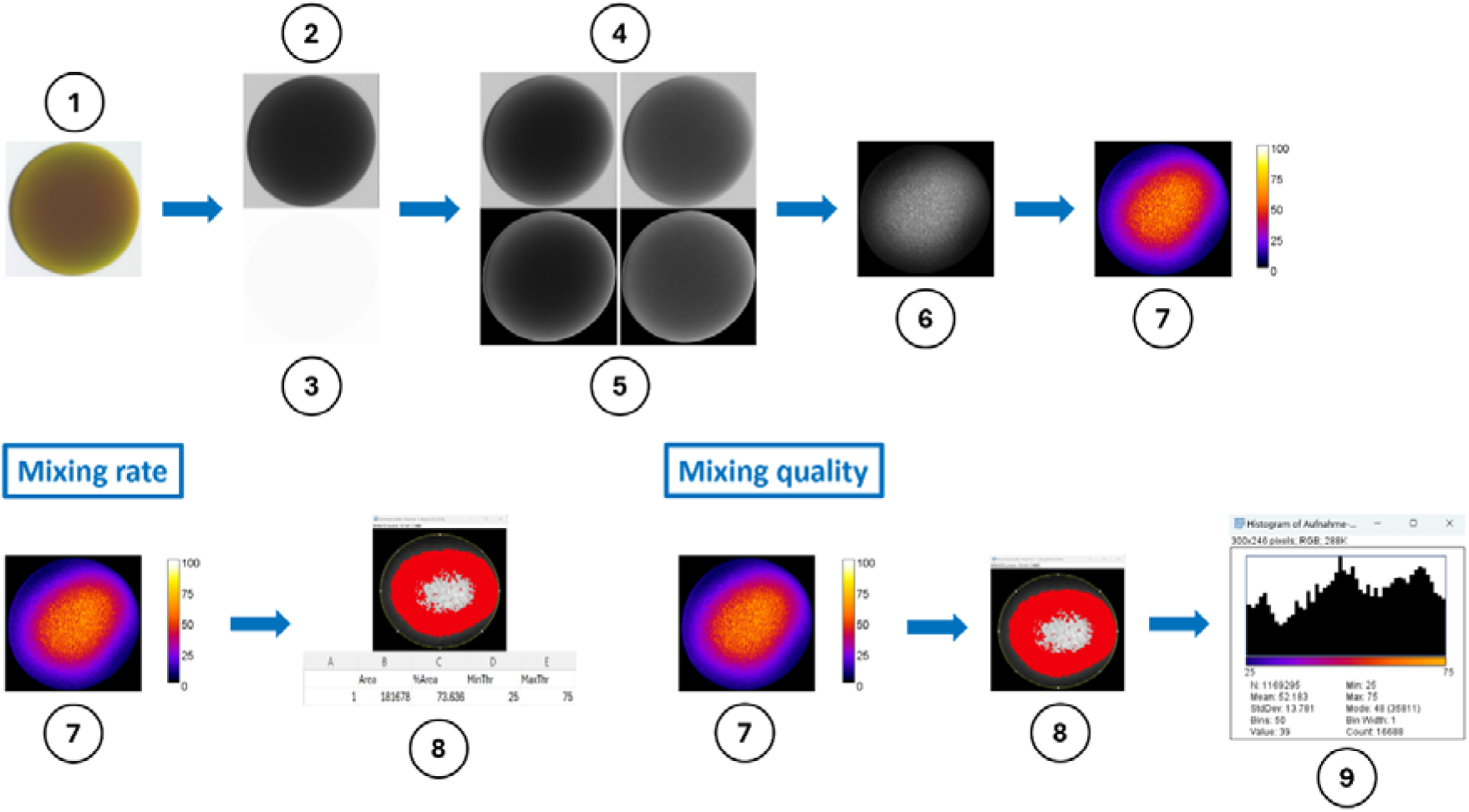
Evaluation of the mixing rate and the mixing quality in Fiji. Top row: 1) Image of the original sample in a .tif-format. If necessary, not required areas are cut off and impurities were removed with median-filters. 2) 8-Bit version of the image. 3) Image-mask. 4) Splitting the image from 1) into the green (left) and red (right) channel. 5) Results of the multiplication from 4) with the image-mask. 6) Results of the division of the red trough the green channel from 5). 7) Final results with the pixelwise coloured evaluation of the sample with concerning calibration bar. Bottom row: Left: 8) Selected area of the sample and result of the calculation of the software for the mixing ratio. Right: 8) Selected area of the sample. 9) Histogram of the concerning area from 8) with the amount of all mixing ratios.

**Figure 3:**
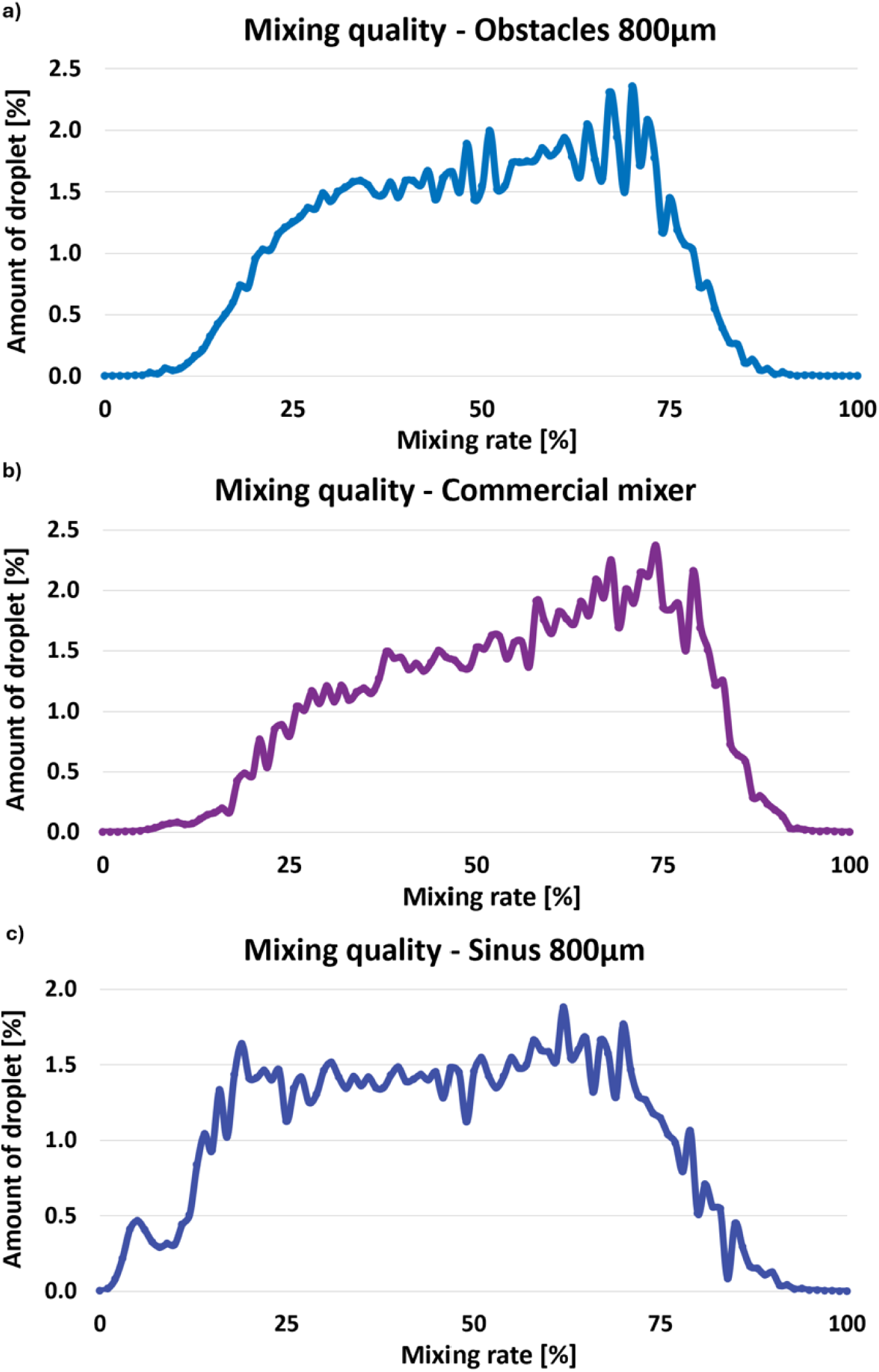
Comparison of the mixing quality for different printheads. a) Evaluation of the mixing quality for a printhead with an obstacle mixing geometry. b) Evaluation of the mixing quality for a printhead with a sinusoidal mixing geometry. c) Evaluation of the mixing quality for a commercial mixing printhead.

**Figure 4:**
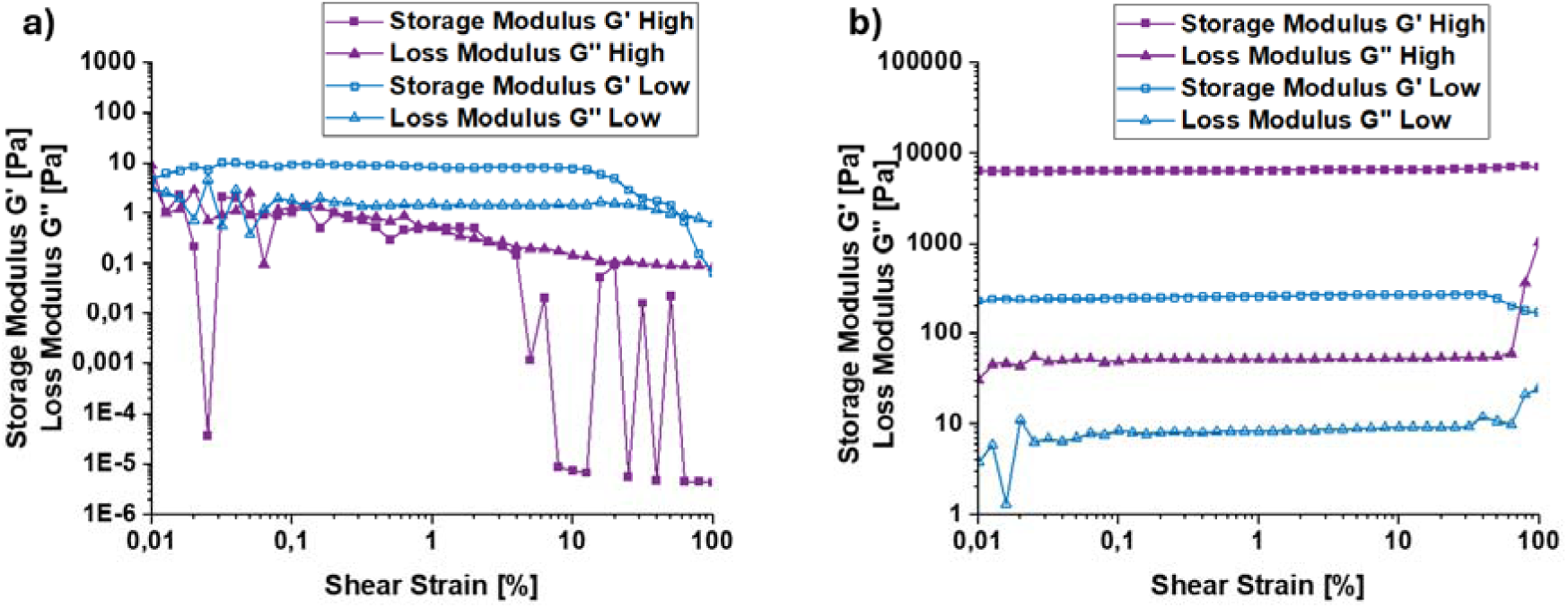
Rheological characterisation of high and low polymer solution before and after UV-crosslinking. a) Amplitude sweep before UV-crosslinking. Until roughly 10 % shear strain the graphs for G’ and G’’ show a parallel progression to the a-axis, which indicates the linear viscoelastic region (LVR). The High variation showed a different behavior. The material left the LVR at a shear strain of 4 %. For small values of the shear strain between 0.01 and 0.1 % the zigzag movement of the graph could be explained due to measurement inaccuracy of the device. Between 0.1 and 4 % shear strain the graph was linear at the chosen frequency. b) Amplitude sweep after UV-crosslinking. Until roughly 40 % shear strain the graphs for G’ and G’’ show a parallel progression to the a-axis for both conditions. Based on these experiments, a value of 1 % shear strain was chosen as amplitude for following rheological tests as both materials were in the LVR during these measurements.

**Figure 5:**
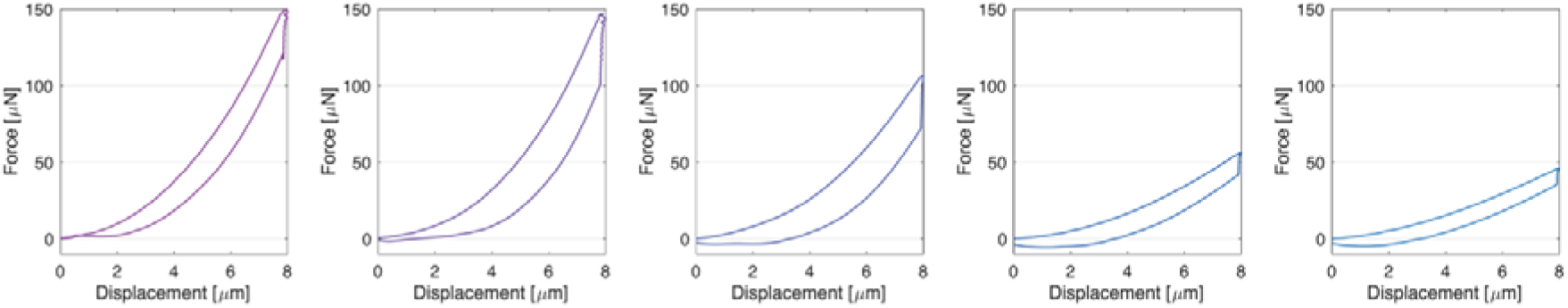
Local mechanical characterisation using nanoindentation of the meander structure displayed in Figure 5b. Exemplary force-displacement curves for all five UV-crosslinked strands of the meander structure.

## Acknowledgements

The authors would like to thank Gizem Karabiyik for her support with the evaluation of the print accuracy and Michael Bartolf-Kopp and Csaba Gergely for their support with taking images of the samples. Further, T.J. thanks the European Union for support on printing strategies (European Fund for Regional Development—EFRE Bayern, Bio3D-Druckproject 20-3400-2-10). K.H., N.S., S.B. and T.J. acknowledge the funding from the Deutsche Forschungsgemeinschaft (DFG, German Research Foundation), Project number 326998133, TRR225 (subproject B09, A07, Z02).

## Notes

### Competing Interest Statement

The authors have declared no competing interest.

